# Mechanism of 2’-Fucosyllactose degradation by Human-Associated *Akkermansia*

**DOI:** 10.1101/2023.10.17.562767

**Authors:** Loren Padilla, Ashwana D. Fricker, Estefani Luna, Biswa Choudhury, Elizabeth R. Hughes, Maria E. Panzetta, Raphael H. Valdivia, Gilberto E. Flores

## Abstract

Among the first microorganisms to colonize the human gut of breastfed infants are bacteria capable of fermenting human milk oligosaccharides (HMOs). One of the most abundant HMOs, 2’-fucosyllactose (2’-FL), may specifically drive bacterial colonization of the intestine. Recently, differential growth has been observed across multiple species of *Akkermansia* on various HMOs including 2’FL. In culture, we found growth of two species, *A. muciniphila* Muc^T^ and *A. biwaensis* CSUN-19, in HMOS corresponded to a decrease in the levels of 2’-FL and an increase in lactose, indicating that the first step in 2’-FL catabolism is the cleavage of fucose. Using phylogenetic analysis and transcriptional profiling, we found that the number and expression of fucosidase genes from two glycoside hydrolase (GH) families, GH29 and GH95, varies between these two species. During mid-log phase growth, the expression of several GH29 genes was increased by 2’-FL in both species, whereas the GH95 genes were induced only in *A. muciniphila*. We further show that one putative fucosidase and a β-galactosidase from *A. biwaensis* are involved in the breakdown of 2’-FL. Our findings indicate that that plasticity of GHs of human associated *Akkermansia* sp. enable access to additional growth substrates present in HMOs, including 2’-FL. Our work highlights the potential for *Akkermansia* to influence the development of the gut microbiota early in life and expands the known metabolic capabilities of this important human symbiont.

**IMPORTANCE:** *Akkermansia* are mucin degrading specialists widely distributed in the human population. *Akkermansia biwaensis* has recently been observed to have enhanced growth relative to other human associated *Akkermansia* on multiple human milk oligosaccharides (HMOs). However, the mechanisms for enhanced growth are not understood. Here, we characterized the phylogenetic diversity and function of select genes involved in growth of *A. biwaensis* on 2’-fucosyllactose (2’-FL), a dominant HMO. Specifically, we demonstrate that two genes in a genomic locus, a putative β-galactosidase and α-fucosidase, are likely responsible for the enhanced growth on 2’-FL. The functional characterization of *A. biwaensis* growth on 2’-FL delineates the significance of a single genomic locus that may facilitate enhanced colonization and functional activity of select *Akkermansia* early in life.

## INTRODUCTION

The foundation of the human gut microbiome begins at birth and is shaped by the initial diet of the newborn, namely human milk (1). Human milk is 87-88% water with the solid fraction made up of carbohydrates (7%), lipids (3.8%), proteins (1%), and other bioactive compounds including hormones and antibodies (2–4). Carbohydrates include lactose and a diversity of other bioactive sugars, called human milk oligosaccharides (HMOs) (5). HMOs are the third most abundant component of human milk yet are indigestible by human enzymes (6, 7), and therefore reach the large intestine largely intact where they act as prebiotics for early bacterial colonizers of the human gut (8). HMOs are composed of only five monosaccharide units (glucose, galactose, N-acetylglucosamine, fucose, and N-acetylneuraminic acid) that form structurally complex linear and branched polysaccharides (9–11). The presence and concentration of these HMOs varies across individuals and is influenced by maternal genetics (12, 13). For example, one of the HMOs, 2’-FL is highly produced by mothers with an active *FUT2* secretor gene, accounting for ∼30% of the total HMOs in these individuals (5). Higher levels of 2’-FL correlate with lower levels of allergies, and lower incidence of eczema and diarrhea in infants (14). In addition to regulating the host immune system, bacterial fermentation of HMOs produces short chain fatty acids (SCFA) that help maintain the intestinal epithelium and regulate appetite (15).

While only a few microorganisms in the infant gut possess the ability to use the entire suite of HMOs, many are capable of fermenting components of HMOs for growth (16–18). For instance, *Bifidobacteria* encode both glycoside hydrolases (GH) and transport systems (19, 20) to catabolize HMOs. However, there are species and strain-to-strain variability in the mechanisms for HMOs degradation across *Bifidobacteria*. For example, *Bifidobacterium longum* subsp. *infantis* uses intracellular enzymes, whereas *Bifidobacterium bifidum* uses extracellular enzymes, pointing to differences in competitive strategies between these organisms (19–22). In addition, other intestinal bacteria such as *Bacteroides thetaiotamicron* and *Bacteroides fragilis* repurpose machinery primarily responsible for the breakdown of mucins to degrade HMOs (23).

The genus *Akkermansia* is comprised of commensal mucin-degrading bacteria that colonize the gastrointestinal tract from early life to adulthood (24) and can also use HMOs for growth (25–27). During growth on HMOs, *Akkermansia* produces acetate, succinate, and propionate (26). The most abundantly produced SCFA, acetate, is involved in fueling gastrointestinal epithelial cells, leading to mucin secretion, and helping to maintain gut barrier integrity(28). In infant guts, there is a positive correlation between the abundance of *Akkermansia* and fucosylated HMOs in mother’s breast milk (29, 30), suggesting a role for this organism in HMO metabolism *in vivo*. Given that *Akkermansia* are largely considered beneficial members of the gut microbiome (31), understanding their ability to degrade HMOs may lead to therapeutic applications.

Although *Akkermansia* are predominantly mucin-degrading specialists, given the structural and compositional similarities between HMOs and mucin oligosaccharides, species may use similar enzyme to break down HMOs (26, 32). Most of the efforts in developing a mechanistic understanding of *Akkermansia* utilization of host glycans thus far have centered around a single isolate of *A. muciniphila* (Muc^T^). The Muc^T^ strain has been shown to use a diversity of GHs including fucosidases, β-hexosaminidases, and β-galactosidases for degrading a variety of human-glycans including HMOs and lactose (26, 32, 33). Recently, ourselves and others have identified and isolated at least four species-level phylogroups (AmI-AmIV) of human associated *Akkermansia*. Thus far, three phylogroups have been named, *A. muciniphila* (AmI), AmII tentatively named *A. massiliensis* (34), and *A. biwaensis* (AmIV, (35)), and the fourth phylogroup, AmIII, is represented by our CSUN-56 isolate and two isolates from Guo and colleagues (36).

We recently demonstrated that all four of these human-associated *Akkermansia* species have the genetic potential and functional ability to metabolize and grow using a variety of HMOs (27). Moreover, some species were more efficient and grew better on individual HMOs. In particular, *A. biwaensis* strain CSUN-19 (AmIV), grew to higher final optical density (OD) on a suite of HMOs and encodes for a larger complement of putative GHs associated with HMO degradation. Interestingly, despite high growth yields when grown on 2’-FL, *A. biwaensis* consumed the least amount of the HMO across *Akkermansia* species. We hypothesized that strains with a greater ability to grow on 2’-FL will have greater expression of GHs or additional GHs required for growth on this substrate. Through phylogenetic analysis, transcriptional profiling, and heterologous expression systems we identified genes encoding putative fucosidases (GH29 and GH95) and β-galactosidase (GH2) from *A. biwaensis* involved in 2’-FL degradation.

## MATERIALS AND METHODS

### Phylogenetic analysis of putative fucosidase encoding genes

To determine the phylogenetic relationships of select glycoside hydrolase (GH) genes across *Akkermansia* species, we analyzed the genomes of 17 publicly available isolates spanning the known phylogenetic diversity of human-associated *Akkermansia* (36–39). Genome accession numbers of isolates used in the analysis are provided in Supplemental Table 1 (Supplemental Table 1). Assembled genomes were submitted to the dbCAN meta server for annotation of GH encoding genes (40, 41). dbCAN uses several tools for annotation and we considered only annotations that agreed using HMMER (42) and DIAMOND (43). All genes annotated as GH2, GH29, and GH95 were identified in the dbCAN output, extracted from the genomes of each strain, imported into MEGA11 (44–46), and aligned using MUSCLE (47, 48). Maximum likelihood trees for each GH family were generated in MEGA11 using default parameters with 100 bootstrap iterations.

### Growth on Fucosylated HMOs

Previously, we showed growth of diverse *Akkermansia* on 2’-FL in medium with 0.5% mucin (27). Since mucin also contains fucose residues with the same α-1,2 configuration as 2’-FL, we first determined the growth of *A. muciniphila* Muc^T^ ATCC BAA-835 (AmI) and *A. biwaensis* CSUN-19 (AmIV) on 2’-FL in the absence of mucin in basal tryptone threonine medium (BTTM) previously described by Ottman and colleagues (25). Briefly, this medium contains 3 mM KH_2_PO_4_, 3.5 mM Na_2_HPO_4_, 5 mM NaCl, 0.5 mM MgCl_2_•6H_2_O, 47.6 mM NaHCO_3_, 0.7 mM CaCl_2_•2H_2_O, 5.6 mM NH_4_Cl, 18 g/L Tryptone (Oxoid, USA), 1.3 g/L L-Threonine, 0.05% Na_2_S-9H_2_O, 1X trace mineral solution (49), and ATCC vitamins (final 1% v/v, manufacturer’s number: ATCC MD-VS, Hampton, NH) with an anaerobic headspace of 70% N_2_ and 30% CO_2_. Stock cultures were maintained in 5 mL BTTM containing 0.4% v/v soluble type III porcine gastric mucin prepared as described previously (27, 49). To obtain mucin-free bacterial cultures, organisms were sub-cultured at 10% inoculum in 5 mL BTTM to which 0.25 mL of 200 mM N-acetylglucosamine (GlcNAc) and 0.125 mL of 200 mM glucose (Glc) was added. Subsequently, for RNAseq, a 10% inoculum was transferred to 10 mL BTTM with the addition of 0.5 mL 200 mM GlcNAc and 0.25 mL of either 200 mM Glc or 200 mM 2’-FL (Glycom, Hørsholm, Denmark). Estimated final concentrations of sugars are 8.3 mM GlcNAc, 4.15 mM Glc, and 4.15 mM 2’-FL. Prior to inoculation, each sugar or mucin was filter sterilized (UNIFLO^TM^ 13mm 0.2mM PES Filter Media, Whatman^TM^) and added to media.

All cultures were incubated at 37°C in a Vinyl Anaerobic Chamber (Coy Laboratory Products, Incorporated, Grass Lake, MI). To monitor growth, 500 uL of each culture was used to determine OD_600nm_ on a Biophotometer plus spectrophotometer (Eppendorf, Biophotometer Plus). Cultures with an OD_600nm_ > 1.0 were diluted with sterile BTTM and re-read. All experimental cultures were grown in quadruplicate.

For follow-up growth experiments with the AkkEH114 (MucT Tn-*Amuc_2072*::*HHJ01_10880* -see below), the following sugar concentrations were used: 4 mM 2’-FL (Glycom, Hørsholm, Denmark) with 0.4% porcine mucin. *Akkermansia* was grown overnight in 5 mL basal mucin medium (27) with 0.4% porcine mucin, sub-cultured in 1 mL BTTM containing media with the appropriate sugars, and 200 uL was transferred to a flat clear-bottom 96-well plate and read hourly on a Spectrostar Nano plate reader (BMG Labtech, USA) for 46 ± 2h with 37°C incubation and 10 seconds of orbital shaking prior to each read. Optical density was converted to OD_600nm_ using a linear equation and the initial (t=0) read was subtracted from all time points as a media blank. All cultures were grown in triplicate and each experiment was repeated three times.

Statistically significant differences between growth at 24 hours was calculated by Tukey’s honest significance multiple comparisons test using R 4.1.1 (R Foundation for Statistical Computing, Vienna, Austria). Comparisons were considered significant if corrected P-values were less than 0.05. Symbol style for figures: nonsignificant (ns), 0.05 (*), 0.01 (**), 0.001(***) and <0.0001(****).

### Targeted Metabolomics

Culture supernatants were collected to measure 2’-FL and Glc degradation during RNA harvest time points. In addition to each parent sugar (2’-FL, Glc, GlcNAc), their individual sugars (fucose, lactose, glucose, and galactose) were also quantitatively measured using high-performance anion exchange chromatography with pulsed amperometric detection (HPAEC-PAD) (50). Culture supernatants were transported frozen (dry ice) where they were thawed in a water bath, vortexed to homogenize, and centrifuged at 7,000 x g for 5 min at 10°C. Then, 0.5 µL of the culture media was injected into the HPAEC-PAD for the detection of the expected analytes described above. Carbohydrate analyses was done as described previously (27).

### RNA Extraction and Purification

RNASeq was performed on cultures grown in BTTM supplemented with GlcNAc and either Glc or 2’-FL (described above) to identify potential genes involved in 2’-FL degradation. Samples were collected at approximately mid-log growth for each conditions/species combination. For *A. muciniphila* Muc^T^, samples were collected after 12h of growth on Glc and 49h for growth on 2’-FL. For *A. biwaensis* CSUN-19, samples were collected after 14h of growth on Glc and 34h for growth on 2’-FL. At the indicated time points, 1 mL of each culture was centrifuged at 4°C at 10,000x g for 7 min. Supernatants from each pelleted sample were removed and placed in the -80°C freezer for metabolomics analysis and pellets were flash frozen in liquid nitrogen for storage in -80°C. RNA was extracted using the phenol/chloroform extraction method outlined in the protocol in RiboPure® Bacteria Kit (Life Technologies Corporation, Carlsbad, CA). Molecular grade Chloroform (MP Biomedicals, LLC, Solon, OH) and Absolute Ethanol (200 proof) (Fisher Bioreagents^TM^, Fisher Scientific, Fair Lawn, NJ) were used for the chloroform/ethanol steps. The elution solution was replaced by water (for RNA work) that is nuclease-free and DEPC-treated (Fisher Bioreagents^TM^, Fisher Scientific, Fair Lawn, NJ). A total of 50µL was eluted in two steps (25µL each step) with ten-minute incubations for each elution step. Two DNase treatments were completed of one hour duration each: DNase I treatment from the RiboPure Bacteria Kit followed by Turbo^TM^ DNase treatment (Life Technologies Corporation, Carlsbad, CA). During Turbo^TM^ DNase treatment, a master mix containing 5µL turbo DNase buffer and 1µL turbo DNase per sample was made prior to adding to the samples. Turbo^TM^ DNase inactivation reagent used was the same one from the RiboPure Bacteria Kit.

RNA was purified and concentrated according to the protocol for RNA Clean and Concentrator Kit - 5 (Zymo Research Corporation, Irvine, CA) with an eluted volume between 30-35µL. Final RNA concentrations were measured using the Qubit^®^ 2.0 Fluorometer with RNA High Sensitivity (HS) Assay Kit (Life Technologies Corporation, Carlsbad, CA) at the completion of the purification. All samples were stored in -20°C until submission to Novogene USA, Inc for Total RNA Sequencing.

### RNA Sequencing and Analysis

RNA samples were sequenced by Novogene USA, Inc using the Illumina^®^ Sequencing HiSeq platform (Illumina, CA, USA). Quality control of RNA samples was assessed using High Sensitivity RNA ScreenTape on TapeStation (Agilent Technologies Inc., CA, USA) and quantified using the Qubit 2.0 RNA High Sensitivity assay (Thermofisher, MA, USA). Ribosomal RNA depletion was performed with Ribo-Zero Plus rRNA Removal Kit (Illumina Inc., CA, USA). Samples were primed and fragmented based on the recommendation of the manufacturer. First strand synthesis was performed with Protoscript II Reverse Transcriptase using a longer extension step of ∼30 min at 42°C. NEBNext^®^ UltraTM II Non-Directional RNA Library Prep Kit for Illumina^®^ (New England BioLabs Inc., MA, USA) was used for the remaining steps of library construction. Final library size was around 350bp with an insert size of 200bp. The final quantity of the library was quantified using the Qubit 2.0 and the quality assessed by the TapeStation D1000 ScreenTape (Agilent Technologies Inc., CA, USA). For the Illumina^®^ HiSeq, 8-nt dual-indices were used and equimolar pooling of libraries was done based on the values from the QC before sequencing with a read length configuration of 150 PE for 20 M PE reads per sample (20M in each direction).

RNA sequencing analysis was performed using the DIY.transcriptomics pipeline (https://diytranscriptomics.com) (51). Raw reads from transcriptomic data were mapped to coding sequences (CDs, no rRNA or tRNA) of each strain using Kallisto (52). The remaining analyses were completed in Rstudio (https://www.rstudio.com) (Supplemental Files 1 and 2). Kallisto transcript abundance measurements were imported into RStudio using the R package *tximport* (53) for filtering and normalization. The Bioconductor/R package *edgeR* (54) *cpm* function was used to create a list of counts per million (cpm) per transcript. Genes with no cpm were filtered out of the dataset using base R’s subsetting method. The *edgeR* function *calcNormFactors* was used to normalize the filtered data using the TMM method (Trimmed Mean of M-values) to the dataset (55).

For analyses of differentially expressed genes (DEGs), a design matrix targeting treatment (Glc versus 2’-FL) was set up to compare cpm of genes in the Glc control versus the 2’-FL treatment. To create a model mean-variance trend and fit the linear model to the data, the *voom* function from the Bioconductor/R package *limma* (56) was used to model a mean-variance relationship, the function *lmfit* was used to fit a linear model to the data, and *makeContrasts* was used to make contrasts of the design matrix between the 2’-FL group versus the Glc group. Bayesian statistics were computed for the linear model fit using the *eBayes* function from *limma* which resulted in moderated statistical values of differential expression (Adjusted P-value, Log fold change, Average Expression, T statistics). To extract DEGs from the linear model fit and Bayesian analysis, the function *decideTests* from *limma* was used to generate a list of significantly regulated DEGs with strict cutoff values (corrected P value ≤ 0.01, log2 fold change ≥ 2) (57).

The R package *EnhancedVolcano* was used to generate volcano plots highlighting the GH29 and GH95 genes for each *Akkermansia* strain with strict cutoff values (p ≤0.05, log2 fold change ≥ 2) (58).

### Cloning and Purification of putative β-galactosidase in E. coli

To characterize one of the β-galactosidases that may be involved in 2’-FL metabolism, the gene fragment of the HHJ01_10865 protein, which encodes the mature GH2 protein lacking the signal peptide (as predicted by SignalP 5.0) (59) was amplified from *A. biwaensis* CSUN-19 genomic DNA using the primers as shown in Supplemental Table 2 (Supplemental Table 2), cut with *Hind*III and *Eco*RI and inserted into the expression vector pET28a(+) (Novagen, Inc.). The resulting recombinant plasmids were transformed into *E. coli* Tuner BL21 for protein overexpression. Bacterial cultures were grown in 50 mL LB medium with kanamycin (50 ug/mL) at 37°C to OD_600nm_ ∼ 0.5, followed by induction with 1 mM isopropyl β-D-1-thiogalactopyranoside (IPTG) for 2 hours at 37°C. Cells were harvested by centrifugation (5,000 g for 5 min at 4°C), the cell pellets were flash-frozen in liquid nitrogen for 5 minutes and stored at -20°C until purification.

Thawed cell pellets were resuspended in 5 mL of purification buffer (50 mM Tris pH 8.0, 200 mM NaCl) and lysed by 4 rounds of sonication [2 minutes of 1 sec on / 2 sec off] with 40% amplitude (Qsonica, USA). Lysates were centrifuged for 30 min at 38,000 g and 4°C and cleared supernatants loaded onto Ni-NTA columns with 0.5 mL bed volume (1018244 Qiagen, Germany). The columns were washed sequentially with 3 column volumes (CV) each of the purification buffer, a low-imidazole wash, and a mid-imidazole wash. Bound proteins were eluted with 4 CV of 100 mM imidazole in purification buffer. Fractions containing recombinant protein, as assessed by SDS-PAGE analysis, were combined and dialyzed against the purification buffer, and the final protein concentration was determined using Qubit protein assay kit (#Q33211 Life Technologies Corporation, USA).

### Activity of Purified Proteins

All enzyme activity and inhibition reactions were performed in 50 mM Tris, 200 mM NaCl, pH 8.0, unless otherwise indicated. Reactions involving ONPG were carried out for one hour at 37°C. Enzymatic products of the analog, orthonitrophenol (ONP), was determined by measuring A_405_ nm using a 96-well plate reader Spectromax M5 (Molecular Devices, USA) and monitored every 2.5 or 5 minutes for one hour following addition of substrate. To determine whether the reaction would be inhibited by excess product concentration, the reactions were run using 5ug of protein at the optimized conditions in the presence of 0.125 mM or 12.5 mM D-glucose, D-galactose, or D-lactose. Reactions were performed continuously at 37°C for one hour in a 96-well plate and reading at A_405_ nm every 2.5 minutes.

### Expression of A. biwaensis HHJ01_10880 in A. muciniphila MucT

The gene encoding the putative *A. biwaensis* GH29 fucosidase HHJ01_10880 (clade 4) was introduced into *A. muciniphila* Muc^T^ using a modified mariner transposon (Tn) delivery system (pSAM_Akk) (60). The plasmid was re-engineered to insert genes within the inverted repeats of Tn system and enable expression of genes of interest in *A. muciniphila* under control of the promoter of *Amuc_1505* (RNA pol) promoter. The resulting plasmid (pE_Akk) contains a *cat* resistance cassette and the *Amuc_1505* promoter flanked by *Mariner* transposase recognition sites.

The coding sequence of HHJ01_10880 was amplified by PCR from *A. biwaensis* Akk2750 (100% sequence match to CSUN-19) (38) with Q5 Hot Start High-Fidelity DNA Polymerase (New England Biolabs) and incorporating a C-terminal HA epitope tag. The amplicon was inserted into *Sal*I- and *Fse*I-digested pE-Akk via In-Fusion Snap Assembly (Takara Bio). The correct sequence of pE_Akk_10880 was confirmed Nanopore sequencing (Plasmidsaurus).

pE_Akk_10880 was introduced into *E. coli* S17-1 λ*pir* strain for conjugative transfer into *A. muciniphila* Muc^T^ as outlined by Davey et al, with minor modifications (60). Briefly, the *E. coli* strain carrying pE_Akk_10880 and *A. muciniphila* Muc^T^ were co-incubated ∼16h in a 10:1 under aerobic conditions on synthetic agar plates (3 mM KH_2_PO_4_, 3 mM Na_2_PO_4_, 5.6 mM NH_4_Cl, 1 mM MgCl_2_, 1 mM Na_2_S·9H_2_O, 47 mM NaHCO_3_, 1 mM CaCl_2_, 40 mM HCl, trace elements and vitamins, 0.2% GlcNAc, 0.2% glucose, 16 g l^−1^ soy peptone, 4 g l^−1^ threonine, 1.25% agar (61, 62) with 0.5 g l^-1^ thioglycolate added and lacking the added sugars (GlcNAc and glucose). After conjugation, cells were diluted and transferred into synthetic medium supplemented with 20uM ciprofloxacin and grown anaerobically for 48 hours. Bacterial cells were passaged twice by diluting 1:10 into fresh medium for 24 hours, before spreading onto synthetic agar plates containing additional 0.25% mucin with ciprofloxacin and chloramphenicol (7ug/mL) and incubated at 37°C until colonies were large enough to pick (∼6 days). Four chloramphenicol resistant colonies were picked and mapping of Tn insert locations identified by arbitrary PCR.

Expression of HHJ01_10880 was confirmed via western blot (Supplemental Figure 1). Samples were separated on a 12% SDS-PAGE gel and subsequently blotted on nitrocellulose membranes (Bio-Rad, 0.22 μM) for 1h at 100V. Membrane blots were blocked with phosphate-buffered saline containing 0.2% Tween-20 (v/v, PBS-T) and 5% nonfat milk (w/v, 5% milk) for one hour. Each membrane was incubated with rat anti-HA primary antibody (Sigma 11867423001; clone 3F10, monoclonal from Roche, 1:5000 in 5% milk) overnight at 4 °C. Blots were washed 3x for 5 minutes in PBS-T at room temperature. Licor anti-rat-680 secondary antibody (LICOR IRDye 680LT Goat Anti-Rat IgG (H + L), Highly Cross-Adsorbed; LIC-925-68029) was added for 1 h at room temperature (1:15000). Blots were washed 3x for 5 minutes in PBS-T at room temperature and developed in the Odyssey Fc Imaging System (LI-COR Biosciences). One of the four mutants with the brightest band, strain Akk-EH114 (MucT Tn*Amuc_2072::HHJ01_10880*) was used in comparative growth experiments with 2’-FL as described above.

## RESULTS

### Evolutionary relationship of GHs involved in 2’-FL metabolism

Enzymes that can target 2’-FL include fucosidases that can break off the α1,2-linked fucose from the parent lactose, which can be further metabolized by a β-galactosidase. To identify mechanisms of 2’-FL degradation by human-associated *Akkermansia,* we first characterized the evolutionary relationship of predicted fucosidases belonging to the GH29 and GH95 families and β-galactosidases belonging to the GH2 family across species. A phylogenetic analysis revealed seven well-supported clades for GH29 and three for GH95 (Figure 1). Generally, when each *Akkermansia* species was represented within a clade, strains from the same species clustered into sub-clades showing phylogenetic congruence for each species. An exception to this pattern is for the GH95 tree, where sequences of *A. biwaensis* were quite divergent within clade 3.

**Figure 1.**
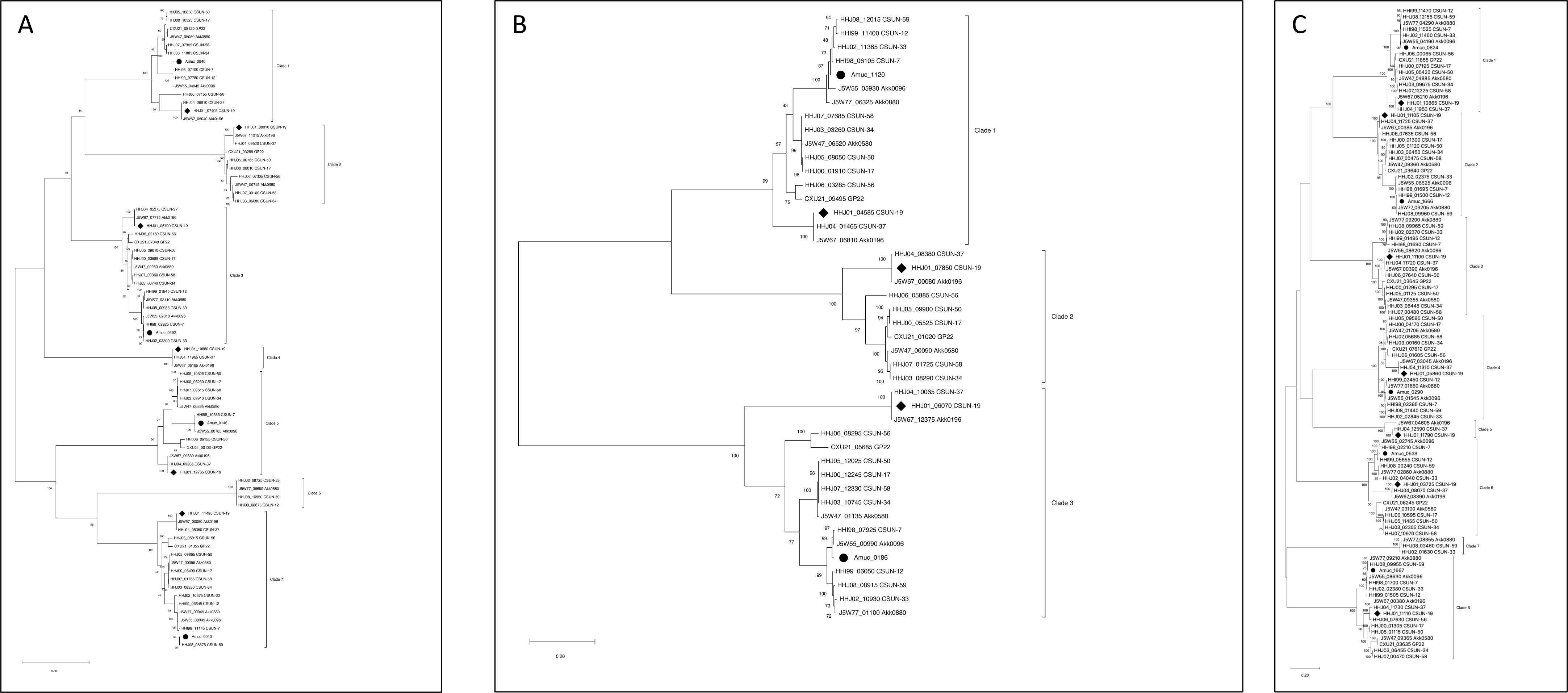
Maximum likelihood trees reveal seven clades of GH29 (A), three clades of GH95 (B), and eight clades of GH2 (C) genes across human associated *Akkermansia* species. Sequences marked by circles represent genes from *A. muciniphila* Muc^T^ (AmI) and sequences marked by diamonds are genes from *A. biwaensis* CSUN-19 (AmIV), the representative strains used throughout this study. All trees were generated using a maximum likelihood heuristic method by executing Neighbor-Join and BioNJ (NJ/BioNJ) algorithms and applied to a matrix of pairwise distances with 500 bootstrap iterations. Trees are drawn to scale, with branch lengths measured in the number of substitutions per site. The trees in (**A**) involved 78 nucleotide sequences and a total of 3,258 positions in the final dataset. For the tree in (**B**), 44 nucleotide sequences and 2,735 sites were analyzed, while in (**C**), 106 nucleotide sequences and 4,919 positions were included. Evolutionary analyses were conducted using MEGA11 (42–44).

Representation of the strains across the clades was not uniform, where some *Akkermansia* species had greater number and diversity of fucosidases. Specifically, across the GH29 genes, *A. biwaensis* strains including strain CSUN-19 used in this study, were present in 6 of the 7 clades including one clade (clade 4) found exclusively in this species (Table 1). In contrast, *A. muciniphila* strains including the type strain Muc^T^, were found in only 4 of 7 clades with none being exclusive to this species. Interestingly, however, clade 6 lacked sequences from either of these two representative species, but contained GH29 genes present in only phylogroup AmIb, a closely related sub-species of *A. muciniphila* previously defined by Becken and colleagues (38).

**Table 1.**
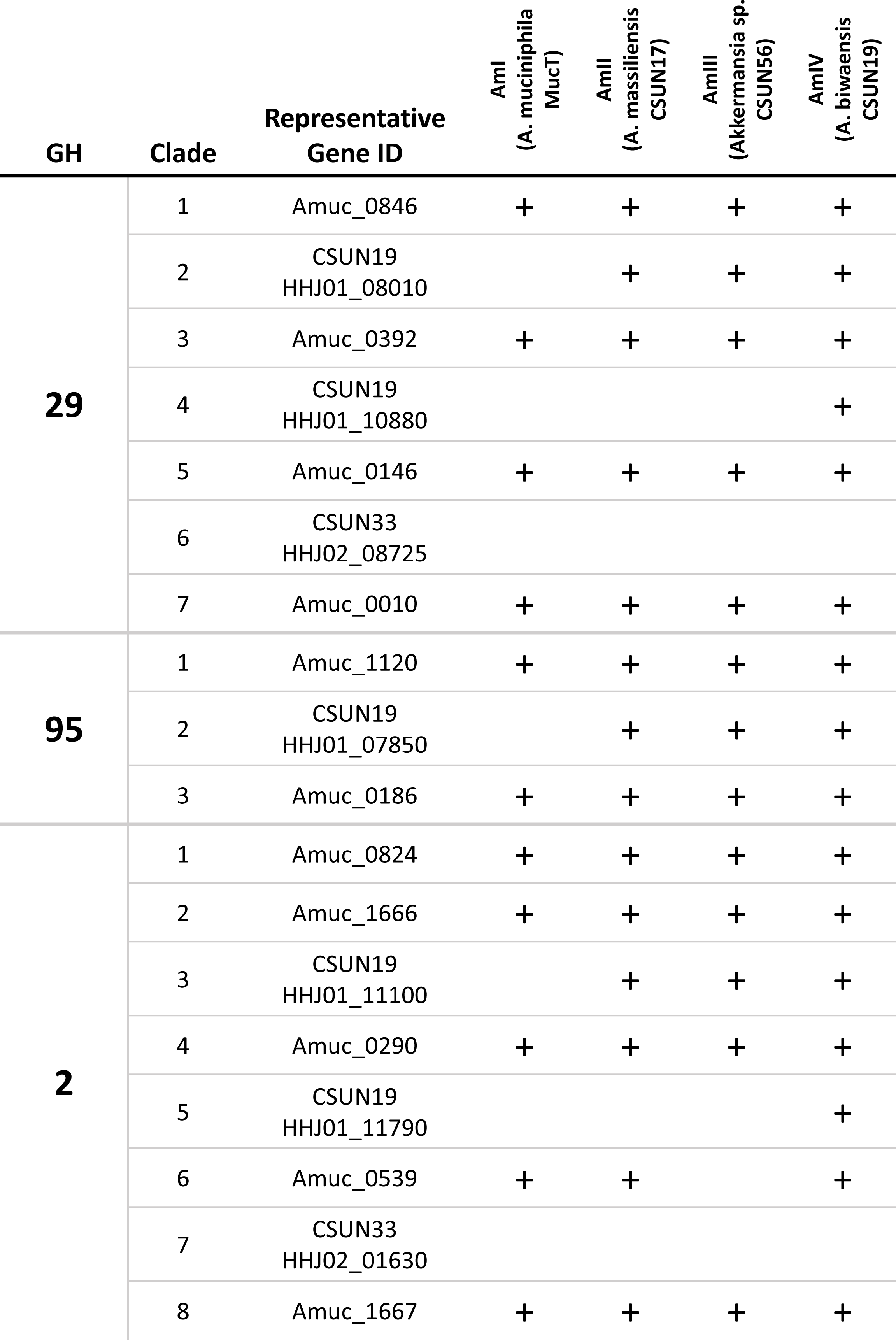
Presence of putative fucosidase (GH29 and GH95) and beta-galactosidase (GH2) genes in representative strains of the four species-level phylogroups reveal a greater suite of predicted fucosidases and beta-galactosidases in phylogroup IV (*A. biwaensis*) of human associated *Akkermansia*. Clade designations are based on maximum likelihood trees (Figure 1).

Similarly, in the GH95 tree, *A. massiliensis*, including strain CSUN-17, *Akkermansia* sp. CSUN-56 from phylogroup AmIII, and *A. biwaensis* including strain CSUN-19 had genes present in all three clades, whereas *A. muciniphila* including the type strain Muc^T^, only had genes in two clades (Figure 1, Table 1).

A similar phylogenetic analysis of GH2 genes in diverse *Akkermansia* revealed eight clades (Figure 1). In line with previous findings, the representative species diverged, in which *A. muciniphila* Muc^T^ had fewer copies of GH2 (5/8 clades) (Table 1), and *A. biwaensis* CSUN-19 had the highest copy number of GH2s in the genome (7/8 clades). Each species also contained unique GH2 sequences; clade 5 was unique to *A. biwaensis* strains while clade 7 was unique to a subset of *A. muciniphila* strains in the AmIb subgroup (Figure 1).

### Growth of A. muciniphila Muc^T^ and A. biwaensis CSUN-19 on 2’-fucosyllactose and glucose

*A. muciniphila* and *A. biwaensis*, the two species with the most divergent number of predicted fucosidases, were used to determine relative growth and generation of sugar metabolites in a minimal media with 2’-FL. Interestingly, although both strains grew using 2’-FL and GlcNAc, the final OD_600nm_ was different across species. By 36 hours the OD_600nm_ of *A. biwaensis* CSUN-19 was nearly double that of *A. muciniphila* Muc^T^ although we are unable to calculate a formal growth rate because of sparse sampling (Figure 2). This contrasts to growth on Glc and GlcNAc, where both species reached similar densities across all time points (Supplemental Figure 2).

**Figure 2.**
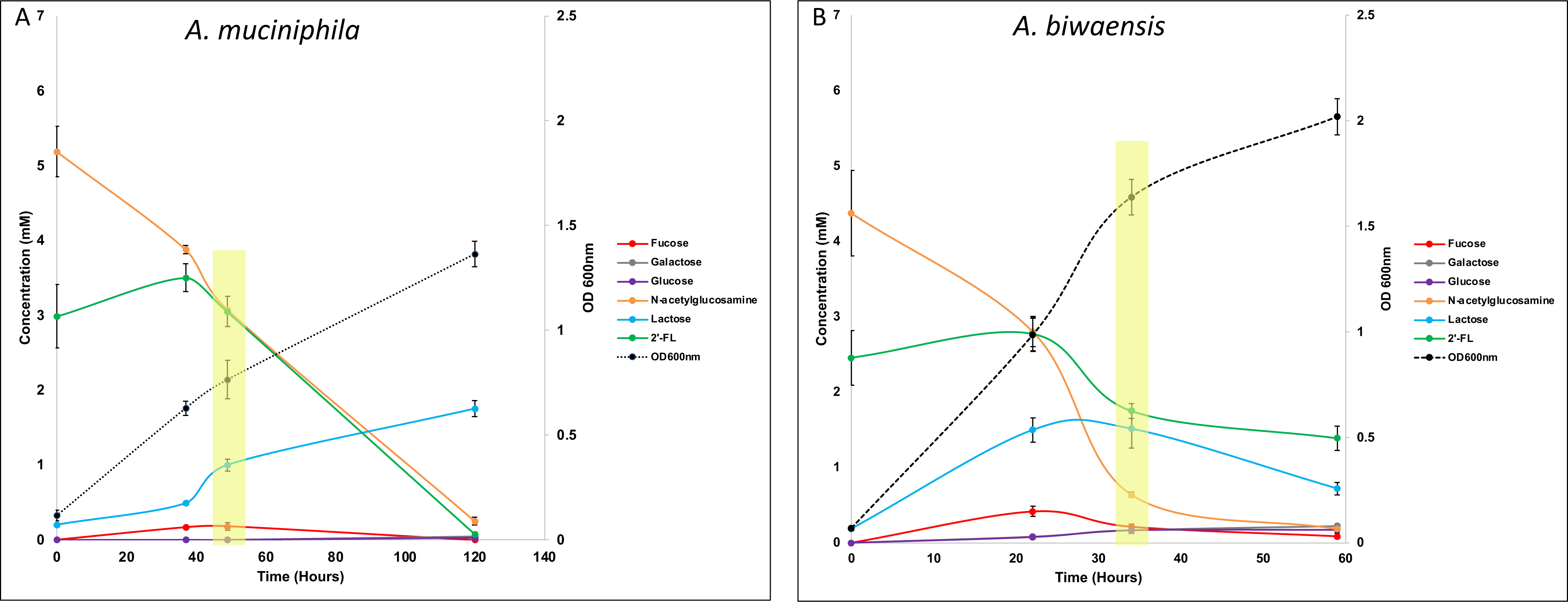
*A. muciniphila* Muc^T^ and *A. biwaensis* CSUN-19 use 2’Fl and GlcNAc for growth. Fucose, lactose, glucose, and galactose, breakdown products of 2’-FL degradation, are detectable in culture media during different stages of growth. Note the differences in time along the y-axis. Asterisks denote time points where cells were collected for RNAseq. Error bars are plus/minus 1 standard deviation of triplicate cultures.

We tested if these species were degrading and consuming the available sugars through similar pathways by measuring the concentrations of the various oligosaccharides (e.g. 2’-FL, lactose) and monosaccharides (e.g. Glc, galactose, GlcNAc). Both strains degraded 2’-FL into fucose and lactose and consumed nearly all GlcNAc by the end of the experiment. Although lactose and fucose were found in the culture medium during growth on 2’-FL, fucose was quickly consumed, whereas lactose accumulated (Figure 2). In addition, small but appreciable amounts of glucose and galactose accumulated in the culture medium of *A. biwaensis* CSUN-19 but not *A. muciniphila* Muc^T^. As anticipated, when grown on glucose and GlcNAc, both strains consumed these sugars within 34 hours of growth (Supplemental Figure 2) (25).

### Gene Regulation of A. muciniphila MucT and A. biwaensis on 2’-fucosyllactose and glucose

We performed an RNASeq analysis of both *A. muciniphila* Muc^T^ and *A. biwaensis* CSUN-19 during mid-log growth on GlcNAc and either Glc or 2’-FL. Because the strains grew at different rates in the different media backgrounds, RNA was extracted from cells harvested during mid-log growth (shading, Figure 2 and Supplemental Figure 2). Importantly, 2’-FL was present in the culture media at the time of sampling, indicating that consumption of this HMO was ongoing. The quality and quantity of all RNA extracts varied but sequencing yields were consistent across samples averaging approximately 15 million reads per sample (Supplemental Table 3). On average, across all samples, 38% of the total reads were aligned to coding sequences (CDs) of each genome. Compared to growth on glucose, 146 of 2,115 (∼6.9%) genes were differentially expressed by *A. muciniphila* Muc^T^ when grown on 2’-FL, of which 51 were down-regulated and 95 were up-regulated (Supplemental Figure 3). Analysis of *A. biwaensis* CSUN-19 identified 615 out of 2,930 (∼21%) differentially expressed genes when grown on 2’-FL versus glucose, of which 344 were down-regulated and 271 were up-regulated.

Initial analysis of the top twenty most differentially expressed genes for each strain revealed limited overlap (Supplemental Tables 4 and 5). The top twenty most differentially expressed genes of *A. muciniphila* Muc^T^ coded for GHs other than GH2, GH29, and GH95, hypothetical proteins, a putative surface layer protein, and two NADPH-dependent oxidoreductases. In contrast, for *A. biwaensis* CSUN-19, none of the top twenty most differentially expressed genes encoded for GHs and constituted mostly hypothetical proteins.

An analysis of the predicted fucosidases, GH29 and GH95 families revealed that in *A. muciniphila* Muc^T^, only Amuc_0846 was significantly upregulated when grown on 2’-FL (log2 fold change >|2|, P_adj_ <0.05), and none of the GH95 genes were significantly regulated (Table 2). For *A. biwaensis* CSUN-19, three GH29 genes were significantly upregulated (log2 fold change >|2|, P_adj_ <0.05), and one GH29 and one GH95 gene were significantly downregulated. The β-galactosidase family GH2 is also likely to be involved in lactose metabolism once fucose is liberated from 2’-FL. Our analysis indicated that two genes, Amuc_1666 and Amuc_0539 were significantly upregulated in *A. muciniphila* Muc^T^ during growth on 2’-FL, whereas GH2 genes in *A. biwaensis* CSUN-19 were not significantly upregulated.

**Table 2.**
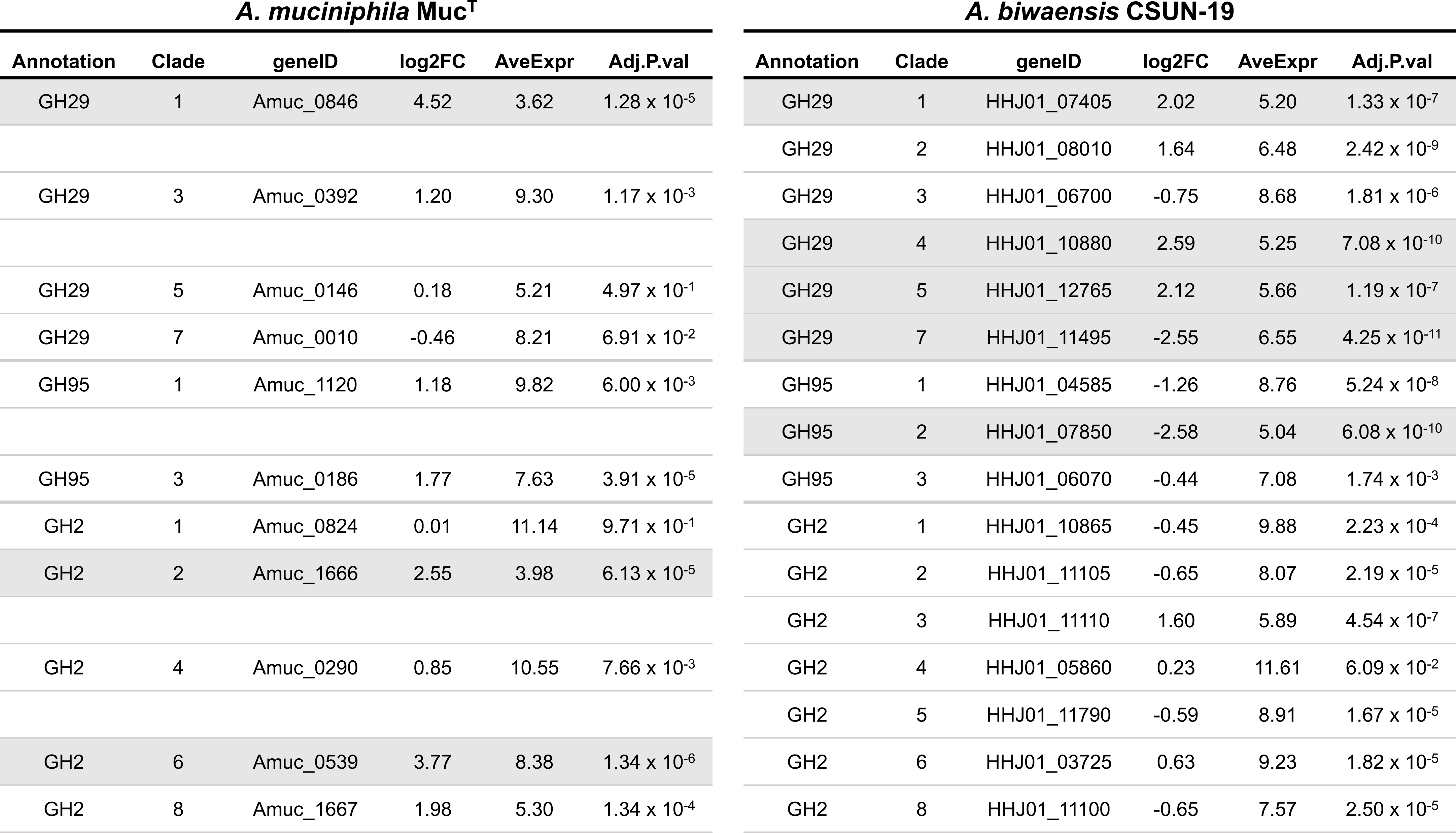
A subset of glycosyl hydrolases are regulated in the presence of 2’-fucosyllactose. RNAseq analysis of the expression of genes encoding glycosyl hydrolases potentially involved in 2’-FL metabolism including fucosidases, such as GH29 and GH95, and β-galactosidases, such as GH2. Fold change (log2FC) in gene expression compared to mid-log growth on glucose for both *A. muciniphila* Muc^T^ and *A. biwaensis* CSUN-19. Average Expression for these genes is based on growth across both conditions. Shaded rows indicates genes that are significantly regulated by 2’-FL.

### Genomic loci of GH2 and GH29/GH95 genes across human-associated Akkermansia

The differences in 2’-FL degradation across species may also be due to genetic variation between the strains, therefore we compared the genomic locus of *A. biwaensis* CSUN-19 containing the most differentially upregulated GH29 (HHJ01_10880) across all four known human-associated *Akkermansia* species (Figure 3). Surprisingly, *A. biwaensis* CSUN-19 was the only species containing this GH29 (HHJ01_10880), agreeing with our phylogenetic analysis. However, all four species have a GH2 ß-galactosidase (*A. muciniphila* Muc^T^ = Amuc_0824; *A. biwaensis* CSUN-19 = HHJ01_10865) within the same genomic locus that may be involved in cleavage of the lactose after removal of the terminal fucose. In addition to the GH2, *A. biwaensis* CSUN-19, *A. massiliensis* CSUN-17, and *Akkermansia* sp. CSUN-56 have genes in this locus encoding a hypothetical protein that is likely a putative sialidase upon further BLASTp search (blue, Figure 3). These same three species that retain the putative sialidase also encode a putative α-galactosidase approximately 7kb upstream (yellow, Figure 3). Interestingly, *A. muciniphila* Muc^T^ is missing all three of these glycoside hydrolases.

**Figure 3.**
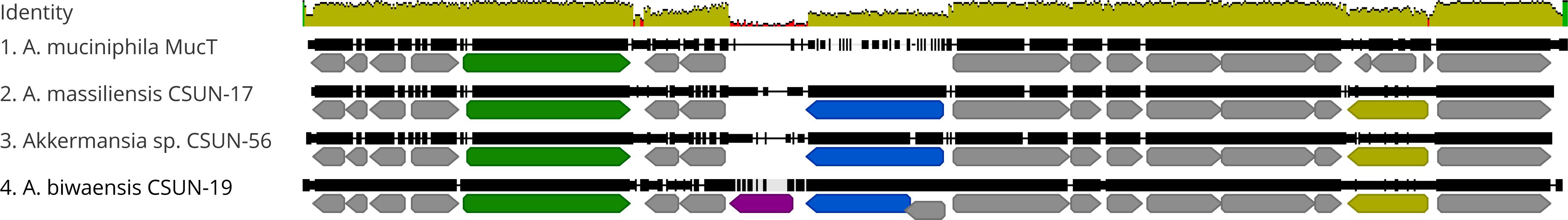
Genomic region containing the FL-2’-regulated GH29 is absent in *Akkermansia muciniphila*. Structure and similarity of the locus containing predicted GH2 and GH29 genes between *A. muciniphila* Muc^T^, *A. massiliensis* CSUN-17, *Akkermansia* sp. CSUN-56, and *A. biwaensis* CSUN-19. Alignment made with Geneious using default parameters. All annotated open reading frames are depicted in grey with the following predicted CAZYmes: GH2 (green), GH29 (purple), GH33 (blue), GH27 (yellow). The region represented is bound by the following genes: Amuc-0820 and Amuc-0836 (Muc^T^), HHJ00_07175 and HHJ00_07250 (CSUN-17), HHJ06_00085 and HHJ06_00010 (CSUN-56), HHJ01_10845 and HHJ01_10930 (CSUN-19).

### Characterization of 2’-FL degrading proteins from A. biwaensis CSUN-19

Next, we aimed to characterize the putative β-galactosidase (HHJ01_10865) and putative α-fucosidase (HHJ01_10880) from *A. biwaensis* CSUN-19 that we expected to be involved in 2’-FL degradation. Towards this, we cloned and purified the putative β-galactosidase, HHJ01_10865 without its signal peptide and determined its enzymatic activity towards ortho-Nitrophenyl-β-galactoside (ONPG). β-galactosidase activity was observed for HHJ01_10865 (Supplemental Figure 4A) in a dose-dependent manner with an optimal pH range of 6.0 to 8.0 and MgCl_2_ dependence (Supplemental Figure 4B,C).

We could not define a α-fucosidase activity for purified HHJ01_10880 on para-nitrophenyl-α-L-fucoside (pNPFuc) or 2’-FL (data not shown). Given the possibility that this GH29 requires secretion or chaperones only present in *Akkermansia* for proper folding, we used a complementary approach of expressing the *A. biwaensis* putative fucosidase in *A. muciniphila*. A transposon-based HHJ01_10880 expression construct was delivered by conjugation into *A. muciniphila* Muc^T^. The insertion sites of the Tn insertions from four clones were confirmed, and one clone (Akk-EH114) was used to assess growth in BTTM + 0.5% mucin supplemented with or without 2’-FL (Figure 4). The Akk-EH114 strain had comparable growth to *A. muciniphila* Muc^T^ in background conditions, but grew significantly better on 2’-FL. The growth pattern of the Akk-EH114 across a 48-hour period is similar to growth of *A. biwaensis* CSUN-19 on 2’-FL, suggesting this enzyme is involved in 2’-FL metabolism (Supplemental Figure 5).

**Figure 4.**
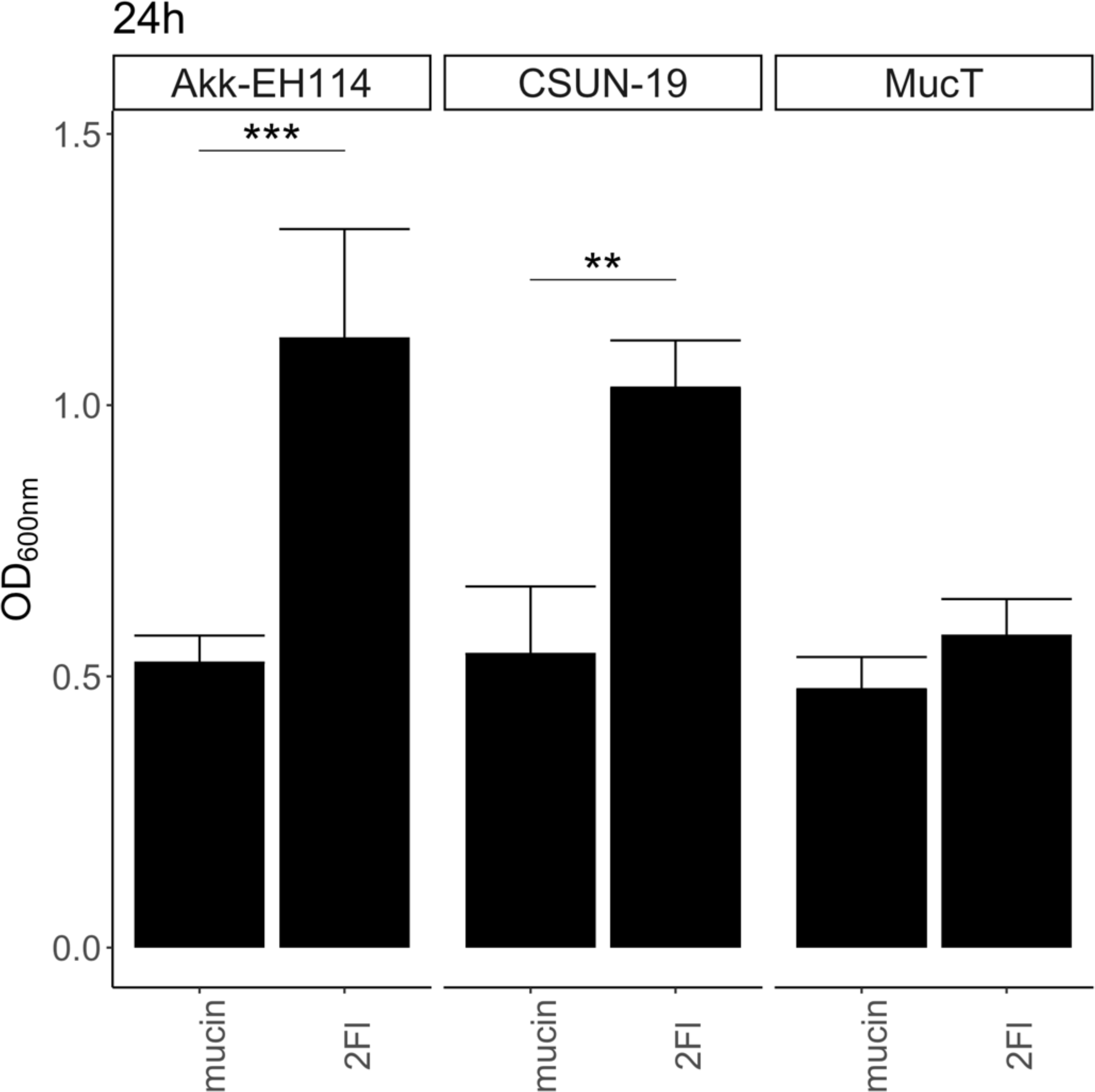
Expression of the *Akkermansia biwaensis* GH29 gene in *A. muciniphila* enhances growth on 2’-FL. *A. muniphila* Muc^T^, A. biwaensis CSUN-19, and *A. muniphila* Akk-EH114 (Muc^T^ expressing HHJ01_10880). Cultures were inoculated into BTTM containing a final concentration of 0.4% mucin and 10 mM 2’-FL. OD_600nm_ was measured at 0h and 24h and normalized to the 0h time point. Error bars represent standard deviation of three biological replicates. Statistically significant differences are calculated by Tukey’s multiple comparisons test with P<0.05. Symbol style: 0.05 (*) and 0.01 (**).

## DISCUSSION

One of the most important functions of HMOs is to facilitate the expansion of beneficial bacteria in the gut microbiota early in life. Recently, different species of human-associated *Akkermansia* have been shown to grow using HMOs (26, 27), expanding their ecological niche and potential importance in the development of the human gut. The mechanisms by which *Akkermansia* use HMOs, specifically the highly abundant 2’-FL, may affect how they interact in and shape both the composition of the infant gut microbiota and development of the infant. Here, we used comparative genomics, phenotypic and transcriptomic profiling, and enzyme activity assays to explore the mechanisms of 2’-FL catabolism in human-associated *Akkermansia* species. We found that strains of *A. biwaensis* possess a higher diversity of both putative fucosidase (GH29 and GH95) and β-galactosidase (GH2) encoding genes than other *Akkermansia* species. Despite this genomic difference, both *A. muciniphila* Muc^T^ and *A. biwaensis* CSUN-19 grew in 2’-FL in media lacking mucin. Furthermore, both strains significantly induced the expression of homologs of a gene encoding a GH29 fucosidase (clade 1), for which the purified protein from *A. muciniphila* has previously been shown to have weak activity against para-nitrophenyl-α-L-fucoside, an analog of 2’-FL (32). Interestingly, a gene unique to *A. biwaensis* strains encoding a different GH29 was also significantly upregulated on 2’-FL. Importantly, expression of this GH29 in *A. muciniphila* was sufficient to enhance its growth to levels similar to *A. biwaensis* in 2’-FL. An additional gene encoding a putative GH2 gene in the same genomic region as the differentially expressed GH29 was confirmed to be a β-galactosidase which is needed to fully deconstruct the 2’-FL trisaccharide. Surprisingly, *A. biwaensis* had a significantly downregulated GH95 fucosidase despite its importance in 2’-FL catabolism. Overall, these findings highlight the different strategies used by *Akkermansia* species to deconstruct fucose-containing HMOs and point to functional differences that may influence the colonization success and health impacts of *Akkermansia* at different host life stages.

### GH29 and GH95 family proteins are involved in 2’-FL catabolism

Analysis of putative fucosidase and β-galactosidase genes across *Akkermansia* species revealed differences in the number and diversity of GH29, GH95, and GH2 genes that was phylogroup specific. Recently, six fucosidases from *A. muciniphila* Muc^T^ have been characterized, demonstrating each enzyme has specific activity on different fucosylated substrates that are diet- or host-derived (32). These corroborate previous work demonstrating that members of the GH95 family hydrolyze 1,2-ɑ-L-fucosidase linkages, while the other dominant fucosidase family, GH29, is divided into two subfamilies in which Group A has little substrate specificity and GH29 Group B hydrolyzes ɑ-1,3/1,4-L-fucosidase linkages (63, 64). Here, we find that *A. massiliensis* CSUN-17, *Akkermansia* sp. CSUN-56, and *A. biwaensis* CSUN-19 contain more copies of both GH29 and GH95 genes than *A. muciniphila* Muc^T^, suggesting capacity to metabolize a broader set of fucosylated substrates or under a broader range of conditions, such as pH or temperature. Several studies have demonstrated optimal performance of fucosidases, sialidases, β-galactosidases, and β-acetylhexosaminidases from multiple organisms, including *Akkermansia*, at specific pH values (26, 64, 65) and fucosidases with different activity depending on the temperature (66). Given that the local pH will vary along the length of the infant intestine, is therefore necessary to understanding the conditions under which these enzymes are expressed to model colonization dynamics (67).

Fucosidase expression patterns observed in *A. muciniphila* Muc^T^ and *A. biwaensis* CSUN-19 under 2’-FL growth conditions suggest that GH29 Group A are the primary enzymes responsible for cleavage of the terminal fucose residue. In our phylogenetic analysis, genes from clade 1 were significantly upregulated in both species. Interestingly, this protein in *A. muciniphila* Muc^T^, Amuc_0846, has weak activity on 2’-FL but strong mucin binding activity (32). Also in agreement, both strains downregulated (significantly only for *A. biwaensis* CSUN-19) another GH29 belonging to clade 7. Previous reports of this purified protein from A. muciniphila Muc^T^, Amuc_0010, demonstrated very weak activity of this purified enzyme against 2’-FL (26, 32). Our results further support these findings suggesting that this fucosidase may not be the primary enzyme for 2’-FL catabolism.

While there were similarities in fucosidase expression patterns across species, there were also notable differences. Most notable was the significant upregulation of HHJ01_10880 in *A. biwaensis* CSUN-19, a predicted GH29 fucosidase absent in other *Akkermansia* species. The purified HHJ01_10880 did not display fucosidase activity using both a substrate analog and the synthetic 2’-FL used in growth experiments. This inability to detect activity is not unexpected, as others have had moderate success rates in obtaining active fucosidases from heterologous expression systems (64, 66).

### β-galactosidases involved in 2’-FL catabolism

Expression levels of the GH2 (HHJ01_10865) located in the same genomic region of *A. biwaensis* CSUN-19 GH29 (HHJ01_10880) was high but not induced by 2’-FL. The activity we observe for recombinant GH2 HHJ01_10865 is similar to other β-galactosidases from HMO-degrading microorganisms (33, 68). This family of enzymes includes proteins with a wide range of activities including β-galactosidase (EC 3.2.1.23), β-mannosidase (EC 3.2.1.25), β-glucuronidase (EC 3.2.1.31), and *exo*-β-glucosaminidase (EC 3.2.1.-) (69). *Akkermansia* harbors other β-galactosidases, such as the GH35 family, that are active on β (1,3)- and β (1,6)-galactoside linkages (70, 71). While the GH35 family is unlikely to target the lactose backbone, HMOs such as Lacto-N-tetraose and Lacto-N-neotetraose, that have a β (1,3) or (1,6) galactoside linkage to N acetyl glucosamine, may be targeted. These linkages are also present in mucin (72), which may implicate β-galactosidases in metabolism of both glycan types. Recently, comparative growth of Tn knockouts in 38 GHs indicated that about 80% of these were not essential for growth on mucin (60). Together with the number, diversity, and shared functions across multiple enzymes, it suggests GH redundancy in *Akkermansia*. Future work using *A. muciniphila* Muc^T^ as a genetic system to express full-length glycoside hydrolases may be used to accurately determine the functional range of these enzymes.

### Genomic linkage of GH genes involved in HMO metabolism

For *Akkermansia* species CSUN-19, the most differentially expressed GH29 when grown on 2’-FL was the predicted fucosidase HHJ01_10880. Interestingly, in the same genomic region as HHJ01_10880, there is also a β-galactosidase 3 genes downstream (Figure 3), albeit on the opposite DNA strand. While this β-galactosidase was not differentially expressed under 2’-FL growth conditions, it was amongst the most highly expressed genes for both organisms (Table 2). In other Gram-negative HMO-degrading bacteria including *Bacteroides*, HMO active GH genes are genomically co-localized and co-regulated in polysaccharide utilization loci (PUL) to efficiently respond to available carbohydrates (23). PULs also contain substrate binding proteins and transport proteins for the target polysaccharides. *Akkermansia* do not have this typical PUL organization despite several GHs map to this region. However, given the regional proximity and high abundance of both transcripts when grown on 2’FL, we propose a model in which the HHJ01_10880 cleaves the fucose by breaking the ɑ1-2 bond, leaving the residual lactose (β1-4) to be cleaved by the adjacent β-galactosidase (HHJ01_10865). Based on the presence of signal peptide sequences, we predict that both the GH29 and the GH2 operating in the periplasmic or extracellular space (26, 60). With this proposed model, after extracellular cleavage of fucose, the lactose would have to be further processed by the β-galactosidase before cellular uptake. Supporting this model, the genome of *A. biwaensis* CSUN-19 lacks an ortholog of the LacY permease, suggesting lactose is not likely imported into the cytoplasm. While this is the first report of introducing and expressing genes in *A. muciniphila* based on the recently developed genetic modification system (60), expansion of this method to include targeted gene knockouts in *biwaensis* is eagerly anticipated to confirm this model. Alternatively, understanding the movement of sugar in these systems would be greatly improved by labeled substrates that can be tracked in real time through the cell.

### Cross-feeding of sugars and metabolites

The difference we observe in metabolic intermediates (i.e., lactose, glucose, and galactose) between *Akkermansia* species may result in differences in how *Akkermansia* supports the growth of other commensal organisms. Like growth in a mucin background, here on synthetic medium, *A. biwaensis* CSUN-19 consumed released lactose whereas *A. muciniphila* Muc^T^ accumulated this disaccharide in the culture medium, which points to potential differences in kinetics of sugar consumption between these species (27). The accumulation of extracellular lactose may be due to secretion of enzymes, as suggested by signal peptides on GHs, observed by us and others (33, 60), which cleave the individual sugars from the backbone before importing the substrates for growth. This could result in cross-feeding potentially leading to support of infant *Bifidobacteria*, many species of which can grow on lactose despite differences in ability to grow on HMOs (22). As an example, cross-feeding interactions have been observed in which *B. breve* can grow on lactose liberated by extracellular breakdown of 2’-FL by *Ruminococcus gnavus* (73). The question therefore remains whether other lactose consumers like *A. breve* will be supported by *Akkermansia* in a species-specific manner.

## CONCLUSIONS

The presence of *Akkermansia* in infants may be correlated with its ability to utilize HMOs. In this study, *Akkermansia* isolates from two divergent species, *A. muciniphila* and *A. biwaensis* were shown to have different growth dynamics on 2’-FL, an abundant HMO in breastmilk. Phylogenetic analysis of putative fucosidases (GH29 and GH95) and β-galactosidases (GH2) involved with the deconstruction of 2’-FL coupled with transcriptional profiling pointed to two genes in a genomic locus that may be responsible for enhanced growth of *A. biwaensis*. We provide evidence that a GH29 present in *A. biwaensis* but absent in *A. muciniphila* is most likely responsible for this phenotype and that both this GH29 and an associated β-galactosidase perform their activities extracellularly (26). In addition to serving as a substrate for *Akkermansia* GH2 enzymes, in a complex ecosystem, the released lactose could also serve as substrate for other members of the infant microbiota. A healthy adult gut microbiome begins in early life where it is shaped by human milk sugars through glycan-degrading microbes. The difference in *Akkermansia* species to break down HMOs suggests a species-specific role in the development of a healthy microbiome. This is significant because it points to the strain-specific mechanisms by which *Akkermansia* may colonize and shape the infant gut, ultimately leading to the rational selection of probiotic bacteria and prebiotic therapies.

## Data availability statement

Data that support the findings of this study are openly available in NCBI BioProject database at SRA, accession number PRJNA1023561.

## Acknowledgements

The authors would like to thank Egle Pakalnyte and Louise Vigsnaes of Glycom for their generous donation of HMOs and their continued support of our work. Thank you to Mike Summers and Melissa Takahashi for providing strains of *E. coli*. Lastly, we would like to thank all current and past members of the Flores lab for the many discussions regarding this project.

## Author Contributions

Conceptualization, G.E.F, L.P. methodology, G.E.F, L.P, A.D.F, E.H. R.H.V.; formal analysis and investigation, G.E.F, A.D.F., L.P., E.L. B.C., visualization, G.E.F, A.D.F., L.P; supervision, G.E.F. and R.H.V. All authors provided critical feedback and helped shape the research, analysis and manuscript. All authors have read and agreed to the published version of the manuscript.

## SUPPLEMENTAL TABLE AND FIGURE LEGENDS

**Supplemental Table 1.** Table of Akkermansia strains used in this study. Accession numbers, phylogroup affiliation, and gene copy number for each strain is given.

| Strain | Accession/BioSample # | Phylogroup | GH2 copies | GH29 copies | GH95 copies | Reference |
| --- | --- | --- | --- | --- | --- | --- |
| *A. muciniphila* Muc<sup>T</sup> | CP001071 | AmI | 5 | 4 | 2 | van Passel *et. al.*, 2011 |
| *A. muciniphila* CSUN-7 | SAMN14614183 | AmI | 6 | 4 | 2 | Luna *et. al.*, 2022 |
| *A. muciniphila* CSUN-12 | SAMN14614184 | AmI | 6 | 4 | 2 | Luna *et. al.*, 2022 |
| *A. massiliensis* CSUN-17 | SAMN14614185 | AmII | 6 | 5 | 3 | Luna *et. al.*, 2022 |
| *A. biwaensis* CSUN-19 | SAMN14614186 | AmIV | 7 | 6 | 3 | Luna *et. al.*, 2022 |
| *A. muciniphila* CSUN-33 | SAMN14614187 | AmI | 7 | 3 | 2 | Luna *et. al.*, 2022 |
| *A. massiliensis* CSUN-34 | SAMN14614188 | AmII | 6 | 5 | 3 | Luna *et. al.*, 2022 |
| *A. biwaensis* CSUN-37 | SAMN14614189 | AmIV | 7 | 6 | 3 | Luna *et. al.*, 2022 |
| *A. massiliensis* CSUN-50 | SAMN14614190 | AmII | 6 | 5 | 3 | Luna *et. al.*, 2022 |
| *Akkermansia* sp. CSUN-56 | SAMN14614191 | AmIII | 5 | 5 | 3 | Luna *et. al.*, 2022 |
| *A. massiliensis* CSUN-58 | SAMN14614192 | AmII | 6 | 5 | 3 | Luna *et. al.*, 2022 |
| *A. muciniphila* CSUN-59 | SAMN14614193 | AmI | 7 | 3 | 2 | Luna *et. al.*, 2022 |
| *Akkermansia* sp. GP22 | SAMN08162556 | AmIII | 6 | 5 | 3 | Guo *et. al.*, 2017 |
| *A. muciniphila* Akk0096 | SAMN18350232 | AmI | 6 | 4 | 2 | Becken *et. al.*, 2021 |
| *A. muciniphila* Akk0880 | SAMN18350241 | AmI | 7 | 3 | 2 | Becken *et. al.*, 2021 |
| *A. massiliensis* Akk0580 | SAMN18350240 | AmII | 6 | 5 | 3 | Becken *et. al.*, 2021 |
| *A. biwaensis* Akk0196 | SAMN18350233 | AmIV | 7 | 6 | 3 | Becken *et. al.*, 2021 |

**Supplemental File 1. R markdown file of basic code and outputs produced during analysis of *A. muciniphila* Muc^T^.**

**Supplemental File 1. R markdown file of basic code and outputs produced during analysis of *A. biwaensis* CSUN-19.**

**Supplemental Table 2.**
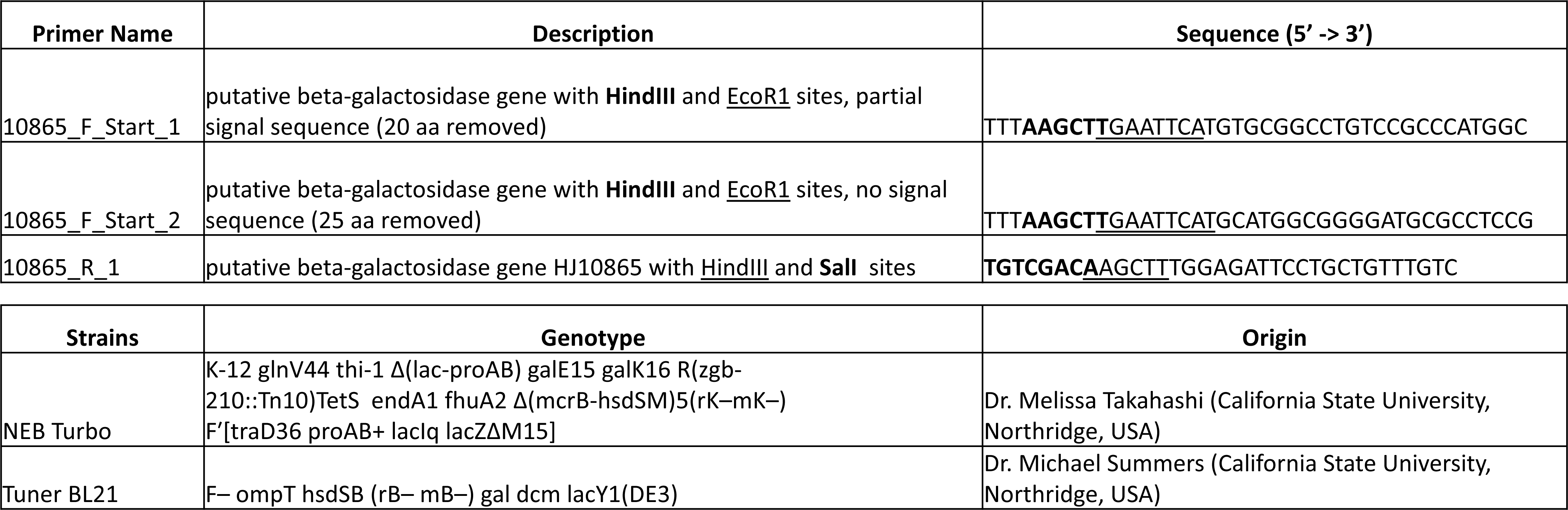
Table of primers and strains used for cloning *E. coli* in this study. Primers used to clone HHJ01_10865 into pET28a for expression in the *E. coli* Tuner strain.

**Supplemental Figure 1.**
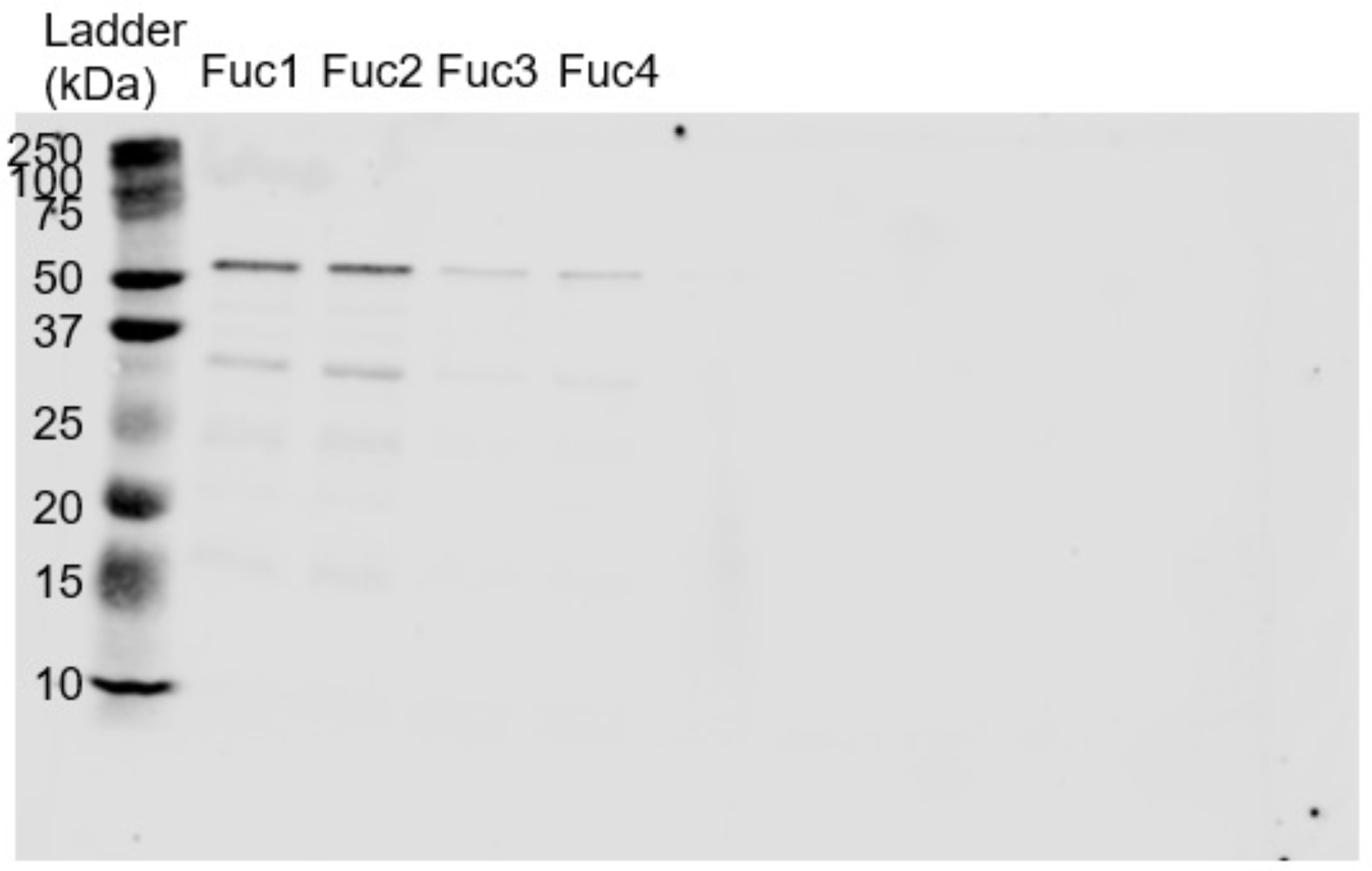
Western blot of four *Akkermansia muciniphila* Muc^T^ strains expression of HA-tagged HHJ01_10880 from *A. biwaensis*. A transposon (Tn) encoding HA-tagged HHJ01_10880 was delivered into *A. muciniphila* Muc^T^ by conjugation and four Cm^R^ colonies characterized by western blot with anti-HA antibodies. Tn insertion sites were identified by inverse PCR and found to occur at *Amuc_1192* (Fuc 1), *Amuc_2072* (Fuc 2), *Amuc_1984* (Fuc 3), and *Amuc_0172* (Fuc 4). Clone #2 (Fuc2) was renamed Akk-EHY114 and used in functional assays.

**Supplemental Figure 2.**
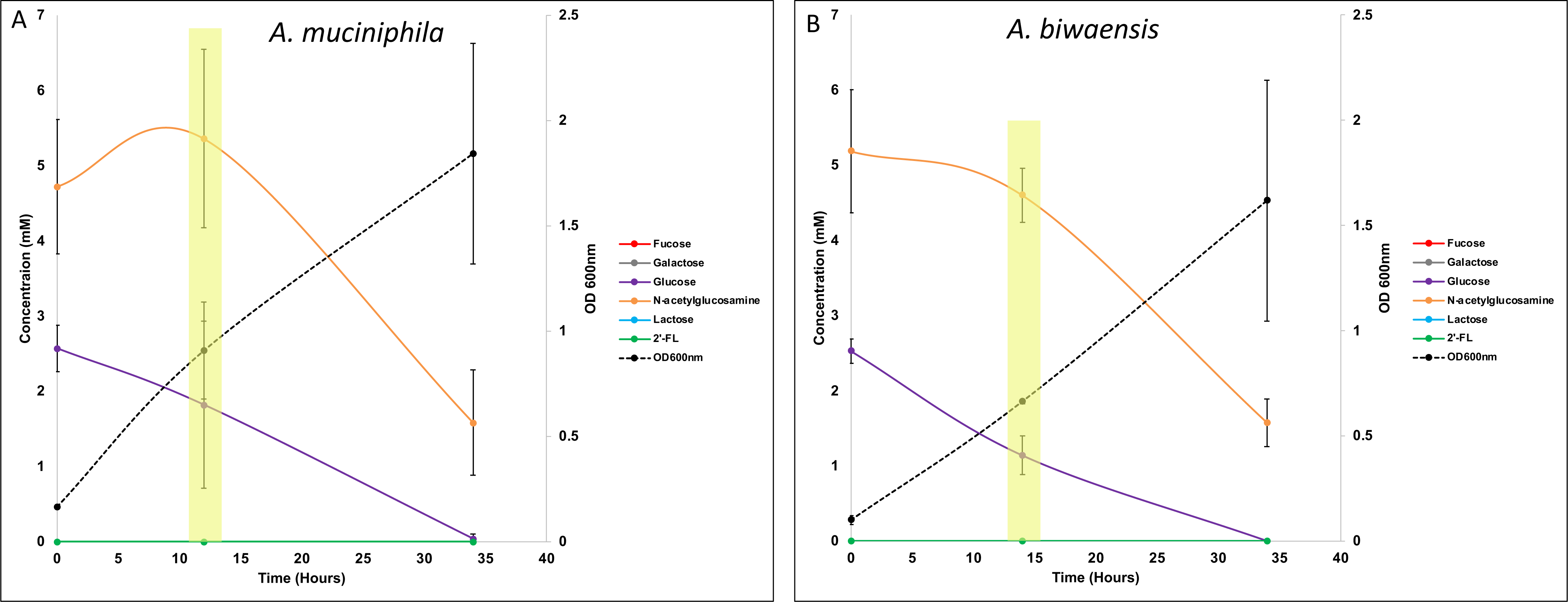
A*k*kermansia species *A. muciniphila* Muc^T^ and *A. biwaensis* CSUN-19 exhibit similar growth kinetics on glucose (Glc) and N-acetylglucosamine (GlcNAc). *A. muciniphila* Muc^T^ (**A**) and *Akkermansia* species CSUN-19 (**B**) degrade Glc (purple line) concomitantly with GlcNAc (orange line) during growth as measured by OD_600nm_ (black dashed line). Areas highlighted in yellow denote time points where cells were collected for RNA extraction and RNAseq. Error bars are plus/minus 1 standard deviation of triplicate cultures.

**Supplemental Table 3.** Whole genome transcriptional profiling resulted in high quality reads matching to the respective genomes. Final measured RNA concentration for each extraction varied within and across samples. Reads were aligned with Kallisto resulting in approximately thirty eight percent non-ribosomal matching to the corresponding coding sequences from each genome.

| Sample ID | Organism | Sugar | Concentration (ng/μL) | Total (ng) | RIN | Q30 | # Reads | Pseudoaligned Reads |
| --- | --- | --- | --- | --- | --- | --- | --- | --- |
| AmucT1 | *A. muciniphila* Muc<sup>T</sup> | Glc | 122.00 | 2,440 | 5.8 | 95.49 | 15,354,070 | 5,715,392 (37%) |
| AmucT2 | *A. muciniphila* Muc<sup>T</sup> | Glc | 41.40 | 828 | 5.6 | 95.46 | 16,183,428 | 6,313,357 (39%) |
| AmucT3 | *A. muciniphila* Muc<sup>T</sup> | Glc | 125.00 | 2,500 | 5.8 | 95.54 | 15,657,002 | 5,763,870 (37%) |
| AmucT4 | *A. muciniphila* Muc<sup>T</sup> | 2'-FL | 9.50 | 190 | 9.8 | 95 | 15,044,641 | 5,211,036 (35%) |
| AmucT5 | *A. muciniphila* Muc<sup>T</sup> | 2'-FL | 10.20 | 204 | 9.6 | 95.12 | 13,766,275 | 4,312,559 (31%) |
| AmucT6 | *A. muciniphila* Muc<sup>T</sup> | 2'-FL | 0.92 | 18 | 9.4 | 92.29 | 15,896,122 | 4,396,982 (28%) |
| CSUN191 | *A. biwaensis* CSUN-19 | Glc | 259.00 | 5,180 | 6.5 | 95.42 | 14,060,510 | 6,482,817 (46%) |
| CSUN192 | *A. biwaensis* CSUN-19 | Glc | 273.00 | 5,460 | 8.3 | 95.48 | 16,576,896 | 7,786,699 (47%) |
| CSUN193 | *A. biwaensis* CSUN-19 | Glc | 291.00 | 5,820 | 6.1 | 95.49 | 14,851,621 | 6,468,715 (44%) |
| CSUN194 | *A. biwaensis* CSUN-19 | 2'-FL | 101.00 | 2,020 | 6.7 | 95.01 | 14,166,132 | 4,951,006 (35%) |
| CSUN195 | *A. biwaensis* CSUN-19 | 2'-FL | 107.00 | 2,140 | 6.5 | 95.17 | 15,492,072 | 5,436,590 (35%) |
| CSUN196 | *A. biwaensis* CSUN-19 | 2'-FL | 78.00 | 1,560 | 6.2 | 95.06 | 15,231,567 | 5,985,852 (39%) |
| CSUN197 | *A. biwaensis* CSUN-19 | 2'-FL | 43.90 | 878 | 5.9 | 95.22 | 14,944,428 | 5,712,869 (38%) |

**Supplemental Figure 3.**
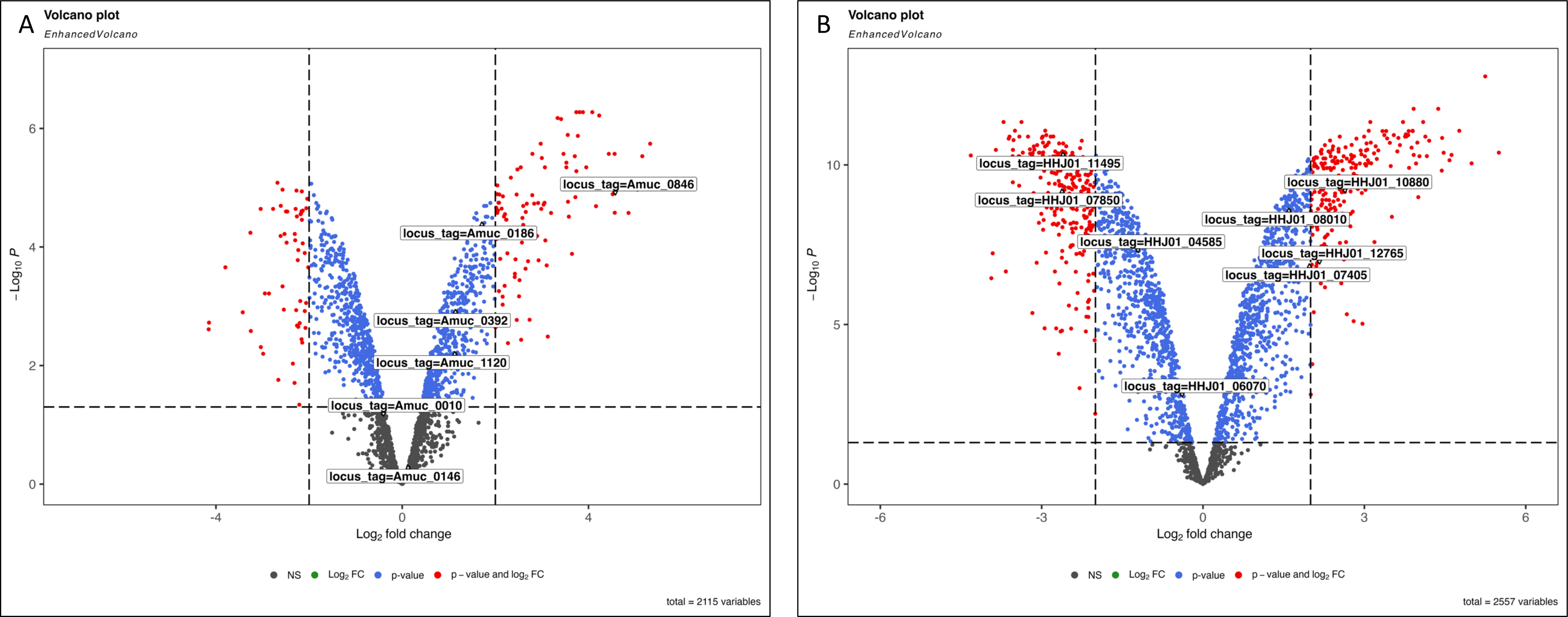
Glycosyl hydrolases genes are regulated during growth in synthetic medium with N-acetylglucosamine (GlcNAc) and 2’-fucosyllactose (2’-FL). Volcano plot of differentially expressed genes of *A. muciniphila* Muc^T^ (**A**) and *A. biwaensis* CSUN-19 (**B**) when grown in synthetic medium with 2’-FL versus glucose. Each dot represents a gene (grey), and colors indicate a P value <0.05 (blue) with a Log2 fold change ≥ 2 (red). Putative fucosidase genes are labeled for *A. muciniphila* (GH29: Amuc_0846, Amuc_0392, Amuc_0010; GH95: Amuc_0186, Amuc_1120) and *A. biwaensis* CSUN-19 (GH29: HHJ01_10880, HHJ01_11495, HHJ01_12765, HHJ01_08010, HHJ01_07405, HHJ01_06700; GH95: HHJ01_07850, HHJ01_04585, HHJ01_06070).

**Supplemental Table 4. Full table of differentially expressed genes for *A. muciniphila* Muc^T^ on 2’-FL vs Glucose.**

**Supplemental Table 5. Full table of differentially expressed genes for *A. biwaensis* CSUN-19 on 2’-FL vs Glucose.**

**Supplemental Figure 4.**
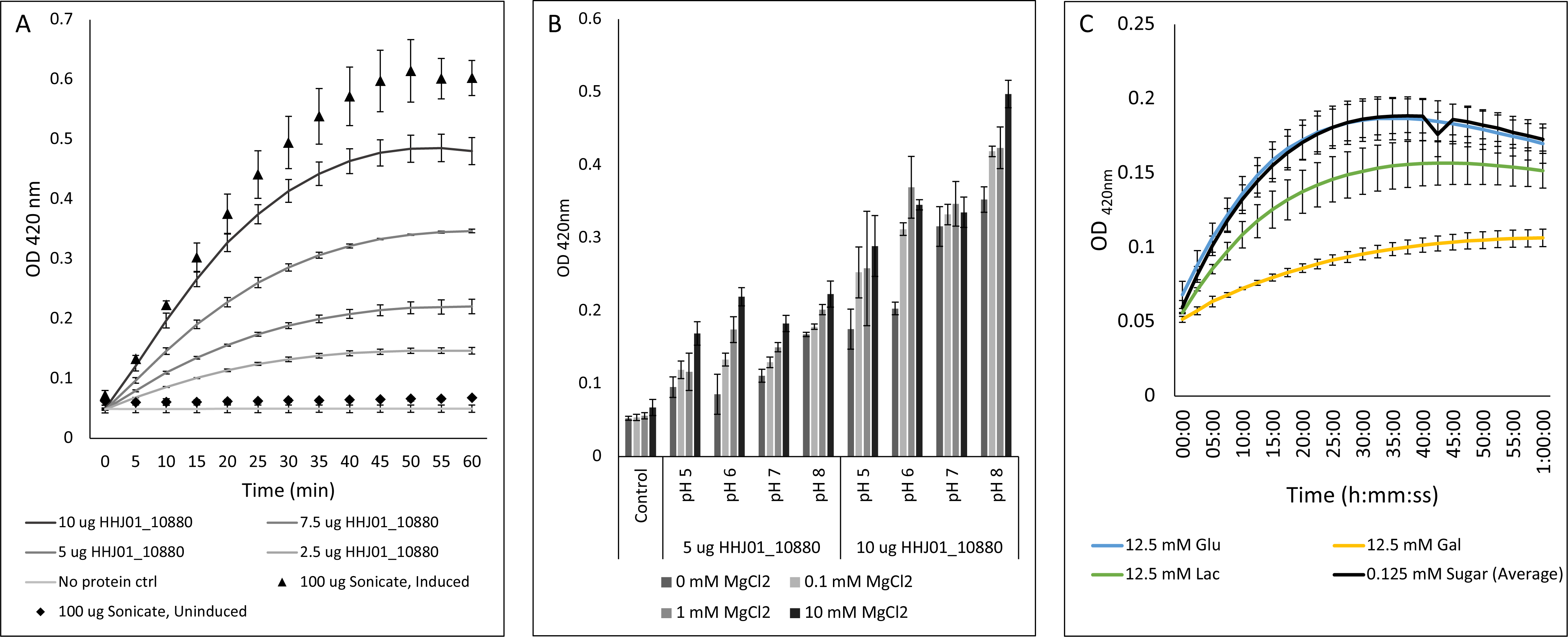
Recombinant GH2 protein from *Akkermansia biwaensis* CSUN-19, HHJ01_10865, has β-galactosidase activity that is strongest at pH 8 with 10 mM MgCl2 and inhibited by galactose. Different concentrations of purified HHJ01_10865 protein or cell lysates from uninduced and induced fractions were incubated at pH8 with 10 mM MgCl2 and ONPG (**A**). Purified protein was incubated at a range of pH and MgCl2 concentrations with ONPG and read at OD_420nm_ after one hour (**B**). 5 ug/mL protein was incubated with ONPG in addition to a 0.05x or 5x concentration of sugars and read every 2 and a half minutes over the course of an hour at OD_420nm_ (**C**). Error bars represent standard deviation of three technical replicates.

**Supplemental Figure 5.**
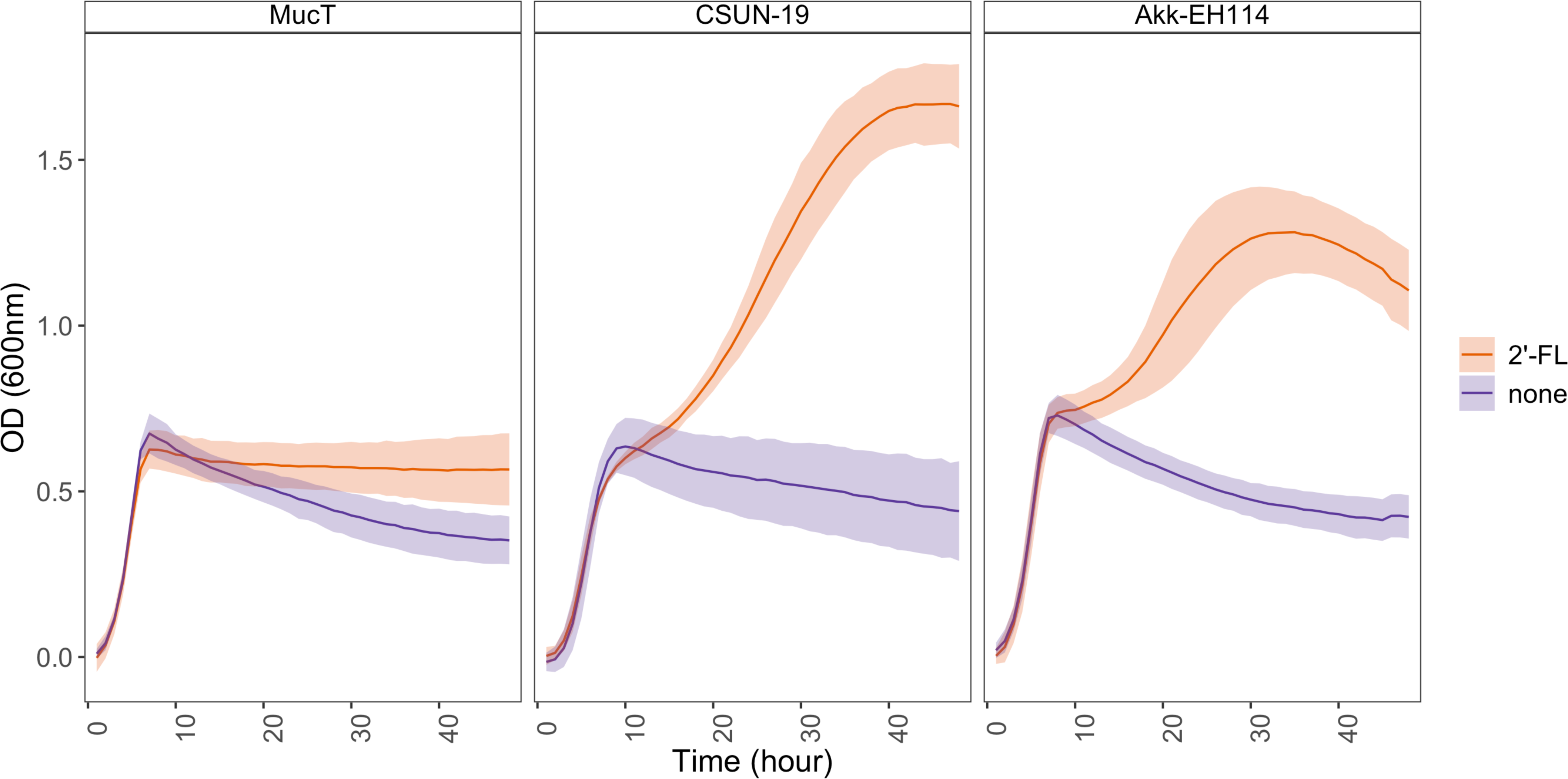
The *Akkermansia muciniphila* strain Akk-EH114 containing the GH29 from *A. biwaensis* CSUN-19 (HHJ01_10880) displays increased growth on 2’Fl. Cultures were inoculated into BTTM containing a final concentration of 0.4% Mucin and 10 mM 2’-FL. Optical density (OD_600nm_) was read every hour for 48h and data was normalized to time zero. Ribbons represent standard deviation of three biological replicates.

